# Structural Basis for pH-Gating of the K^+^ Channel TWIK1 at the Selectivity Filter

**DOI:** 10.1101/2021.11.09.467928

**Authors:** Toby S. Turney, Vivian Li, Stephen G. Brohawn

## Abstract

TWIK1 is a widely expressed pH-gated two-pore domain K^+^ channel (K2P) that contributes to cardiac rhythm generation and insulin release from pancreatic beta cells. TWIK1 displays unique properties among K2Ps including low basal activity and inhibition by extracellular protons through incompletely understood mechanisms. Here, we present cryo-EM structures of TWIK1 in lipid nanodiscs at high and low pH that reveal a novel gating mechanism at the K^+^ selectivity filter. At high pH, TWIK1 adopts an open conformation. At low pH, protonation of an extracellular histidine results in a cascade of conformational changes that close the channel by sealing the top of the selectivity filter, displacing the helical cap to block extracellular ion access pathways, and opening gaps for lipid block of the intracellular cavity. These data provide a mechanistic understanding for extracellular pH-gating of TWIK1 and show how diverse mechanisms have evolved to gate the selectivity filter of K^+^ channels.

## Introduction

Two-pore domain K^+^ channels (K2Ps) generate leak-type currents regulated by diverse chemical and physical stimuli including temperature, membrane tension, signaling lipids, and pH to control the resting membrane potential of cells (Enyedi and Czirjak, 2010). K2Ps assemble as homo- or hetero-dimers, with each subunit containing four transmembrane-spanning helices (TM1-TM4), two selectivity filters (SF1-SF2), two pore helices (P1-P2), and two helices that comprise an extracellular cap (EC1-EC2) (Enyedi and Czirjak, 2010; Natale et al, 2021). TWIK1 (**T**andem of Pore Domains in a **W**eak **I**nward Rectifying **K**^+^ Channel, also known as K2P1 or KCNK1) was the first K2P to be discovered and is widely expressed, including at high levels in the brain and heart in humans (Lesage et al, 1996).

TWIK1 is a pH-gated K^+^ channel that responds to extracellular acidification. TWIK1 is predominantly closed at pH 5.5 and open at pH 7.5 with a midpoint pH ≈ 6.7 (Rajan et al, 2005). H122 has been identified as the extracellular proton sensor and is located directly above the selectivity filter. The TWIK1 selectivity filter is unique among K^+^ channels, having diverged from the highly conserved T(L/V/I)G(Y/F)G motif to T**T**GYG for SF1 and TIG**L**G for SF2 (Chatelain et al, 2012). Under conditions of low pH and low [K^+^]_ext_ (< 5 mM), TWIK1 reversibly enters a non-selective conductive state rather than closes as seen at higher [K^+^]_ext_ (Ma et al, 2011; Ma et al, 2012; Chatelain et al 2012). This pH- and [K^+^]_ext_-dependent selectivity loss is postulated to underlie the seemingly paradoxical hyperpolarization of kidney and pancreatic β cells upon TWIK1 deletion(Millar et al, 2006; Chatelain et al, 2012) and TWIK1-dependent depolarization of human cardiomyocytes under hypokalemic conditions (Ma et al, 2011; Ma et al, 2012).

Currently, structural insight into TWIK1 is limited to a crystal structure determined at pH 8 (Miller et al, 2012). Consistent with the channel being open at high pH, the TWIK1 selectivity filter adopted a conductive conformation with K^+^ ions bound at canonical sites S0-S4. However, the ion conduction path was blocked in the intracellular cavity by a bound detergent or lipid acyl chain through a lateral membrane opening between transmembrane helices TM2 and TM4. Analogous acyl chain block has been observed in other K2P structures determined in detergent (Brohawn et al, 2014; Dong et al, 2015) or lipid nanodiscs (Li et al, 2020) and hydrophobic drugs that inhibit K2Ps, including the antidepressant norfluoxetine, occupy a similar site (Dong et al, 2015; Rodstrom et al, 2020). How exactly this site is involved in TWIK1 gating is unclear (add Aryal et al, 2014; Aryal et al, 2015). Mutational, spectroscopic, and electrophysiological evidence rather implicate the TWIK1 selectivity filter in gating conformational changes in ways that are distinct from other K^+^ channels (Ma et al, 2011; Chatelain et al, 2012; Tsukamato et al, 2018; Nemation-Ardestani et al, 2020). Still, the structural and mechanistic basis for TWIK1 pH-gating remains unknown.

## Results and Discussion

We found that full-length *Rattus norvegicus* TWIK1, which shares 96% sequence identity to human TWIK1, was well-expressed in *Pichia pastoris* and biochemically suitable for structural studies. Like human TWIK1 (Rajan et al, 2005; Ma et al, 2011; Chatelain et al 2012), rat TWIK1 was inhibited by low pH_ext_ in high [K^+^]_ext_ (96 mM) (Fig. 1A), and displayed a linear current-voltage relationship. In low [K^+^]_ext_, K^+^ selectivity was reduced upon exposure to low pH_ext_ (2 mM K^+^) (Fig. 1B), and the channel displayed inward rectification at low [K^+^]_ext_ and high pH_ext_ (Lesage et al 1996). To eliminate intracellular sequestration, preclude possible SUMOylation, and increase functional expression of TWIK1(Rajan et al, 2005; Feliciangeli et al, 2007; Feliciangeli et al, 2010), all electrophysiological recordings were conducted in a triple mutant background (TWIK1 K274E,I293A,I294A), similarly to previously described studies (Chatelain et al, 2012).

**Figure 1.**
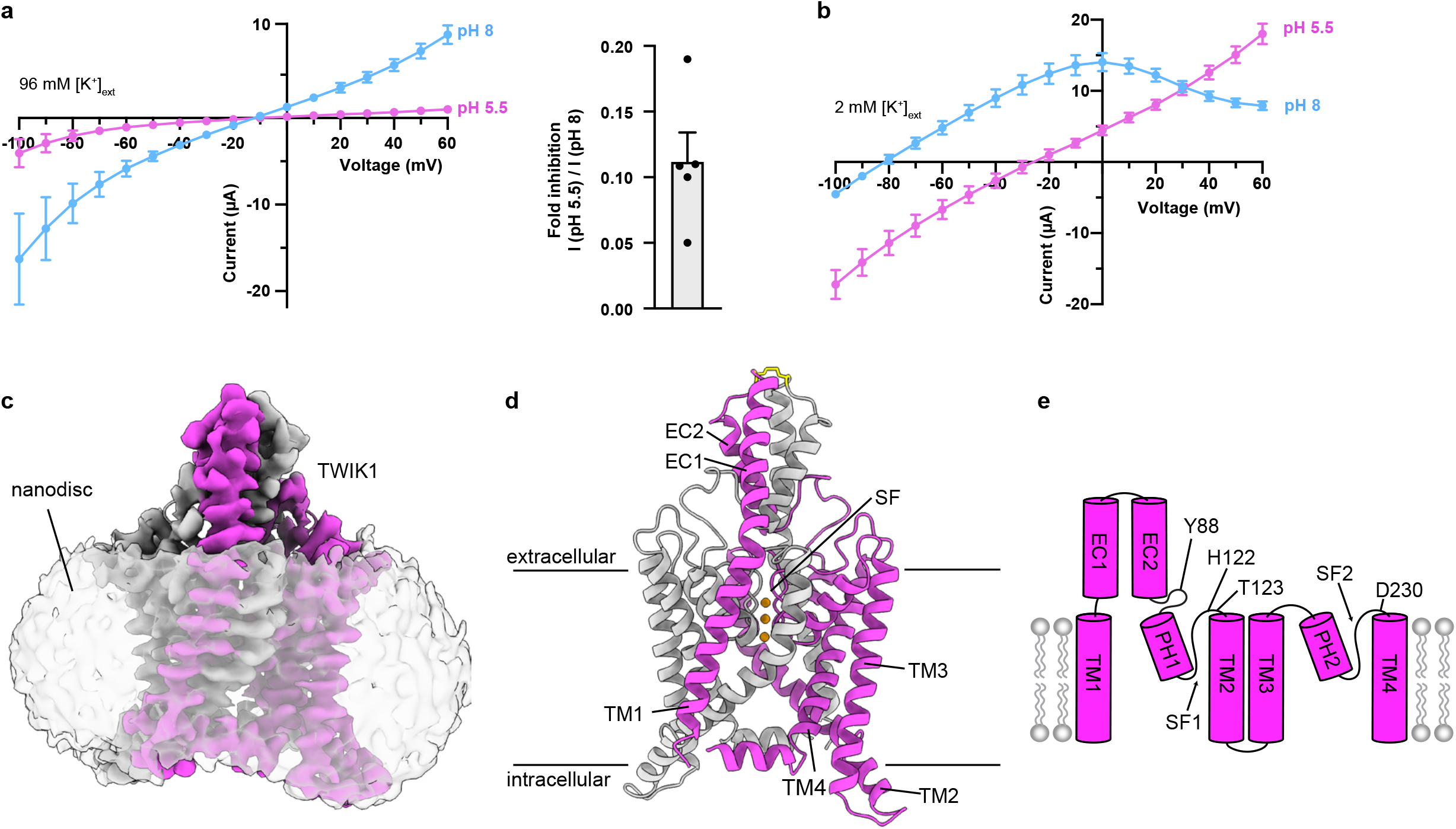
Structure and function of TWIK1. **(a**,**b)** Current-voltage relationships from TWIK1-expresing cells at high and low pH_ext_ in **(a)** high [K^+^]_ext_ and **(b)** low [K^+^]_ext_. (**a, right**) Fold inhibition in high [K^+^]_ext_ (I_pH 5.5_ / I_pH 8_ at 60 mV) = 0.11 ± 0.02. Currents are mean ± s.e.m. from n=5 cells. **(c)** Cryo-EM map at pH 5.5 viewed from the membrane plane. Nanodisc is transparent and TWIK1 subunits are magenta and white. **(d)** TWIK1 model colored as in **(c)** with K^+^ ions orange and disulfide yellow. **(e)** Cartoon representation of a TWIK1 protomer with four transmembrane helices (TM1-TM4), two extracellular cap helices (EC1-EC2), two pore helices (PH1-PH2), two selectivity filters (SF1-SF2), and key residues involved in extracellular pH-gating indicated.

To capture structures of TWIK1 in a lipid environment, we reconstituted the channel in nanodiscs containing the lipids DOPE, POPC, and POPS (Fig. S1) and determined its structure by cryo-EM at pH 7.4 and pH 5.5 to approximately 3.4 Å resolution (Fig. 1C-1E, Fig. S2-S4, Table S1). The channel is two-fold symmetric at both pH values and enforcing C_2_ symmetry of the reconstructions improved map quality. Amino acids W20 to Y281 are clearly defined and modeled in the low pH structure, corresponding to a visible mass of 59 kDa. The high pH reconstruction is more anisotropic due to preferred orientations of particles (Fig. S4A,S4D). Some regions, particularly the intracellular termini and TM2-TM3 linker, are less well resolved in the high pH map, but amino acids G24 to F280 could still be confidently modeled. TWIK1, like other K2Ps, is a domain-swapped homodimer (Fig 1D). The initial crystal structure of TWIK1 (Miller 2012) was modeled without a domain swap likely due to poor local resolution of the distal cap region, as was the case for the related K2P TRAAK (Brohawn et al, 2012; Brohawn et al, 2013).

Consistent with functional data, TWIK1 adopts an open conformation at pH 7.4 and a closed conformation at pH 5.5. The high pH structure shows an unobstructed path from the intracellular to extracellular solutions through the channel cavity, selectivity filter, and bifurcated extracellular tunnels under the helical cap (Fig. 2A,2C-D). At low pH, gating conformational changes seal the top of the selectivity filter and extracellular pathways under the helical cap, forming constrictions with radii of ∼0.5 Å and ∼0.8 Å that are too small for K^+^ ions to pass (Fig. 2B-2D). The conduction path is further blocked at low pH at the intracellular side of the selectivity filter by lipid acyl chains bound within the channel cavity (Fig. 2B, 2D).

**Figure 2.**
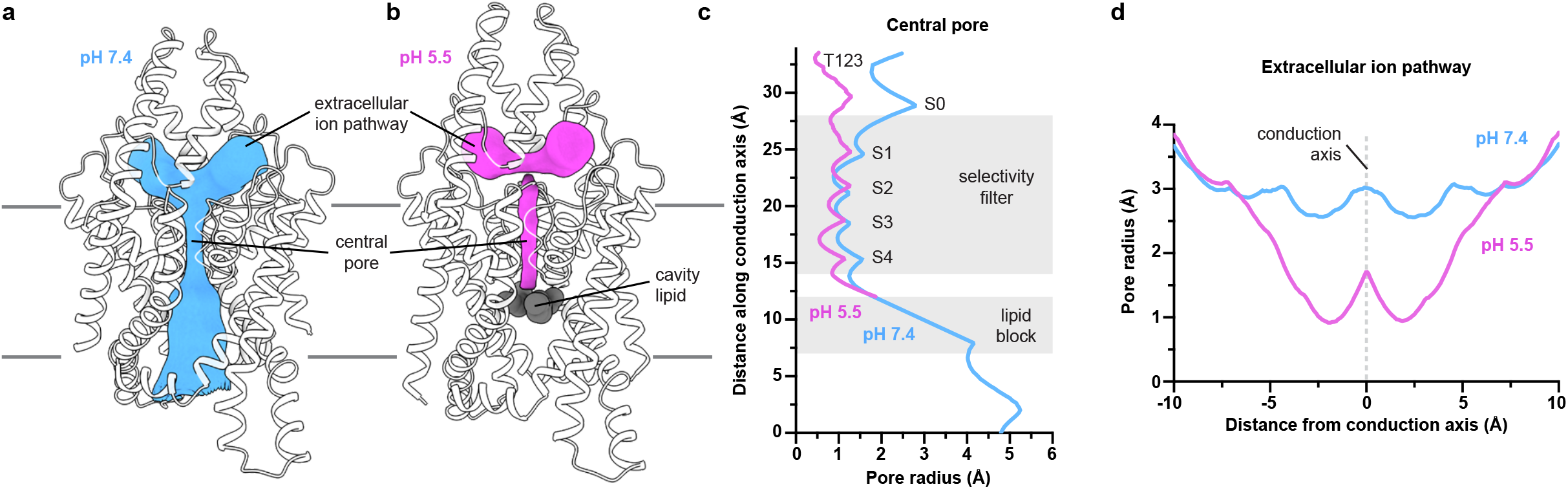
TWIK1 conduction pathways at high and low pH. **(a**,**b)** Structures of TWIK1 at **(a)** pH 7.4 and **(b)** pH 5.5 with the surface of conduction pathways shown in grey. **(c)** Radius of the central pore as a function of distance along the conduction pathway. Positions of the selectivity filter, lipid block at low pH, K^+^ coordination sites S0-S4, and T123 are indicated. **(d)** Radius of the extracellular ion pathways as a function of distance from the central conduction axis.

At high pH, the selectivity filter adopts a nearly four-fold symmetric conformation with K^+^ ions bound at five positions in sites S0-S4, as observed in other conductive K^+^ channels and the previously reported TWIK1 crystal structure at pH 8 (Cα r.m.s.d. within pore helices and selectivity filters = 0.6 Å). The intracellular side of the filter is in a similar conformation at both pH values, but the extracellular side of the filter is dramatically rearranged at low pH. Rotation of Y120 and L228 carbonyls at the top of S1 nearly 180° away from the conduction axis and displacement of the SF1-TM2 and SF2-TM4 linkers disrupts the K^+^ coordination environment at S1 and S0 sites. Due to the change in coordination environment, K^+^ binding sites S0-S1 are unoccupied at low pH and density corresponding to K^+^ ions is only observed at positions S2-S4.

Gating conformational changes are propagated from the top of the selectivity filter (Fig. 3). At high pH, the residues immediately after the selectivity filter sequences, H122 and D230, project back behind the filter and away from the conduction axis. Upon protonation, H122 flips upward towards the helical cap. The adjoining linker region connecting SF1 to TM2 (H122 to D128) kinks outward towards the SF2-TM4 linker on the opposing subunit. D230 and the SF2-TM4 linker flip upward and outward in a similar way, though to a lesser degree, positioning D230 ∼3.1 Å from protonated H122. Movement of the four linkers is concerted because any one linker in a pH 5.5 position sterically clashes with a neighboring linker in a pH 7.4 position. Viewed from above, the linkers push one another into pH 5.5 positions like four dominos falling counterclockwise and form a new set of interactions. D230s form salt bridges with H122s on either side of an interaction between opposing T123s. These three amino acids from each protomer form a zipper at low pH directly above the selectivity filter to block ion conduction and stabilize the non-conductive conformation (Supplemental Video 1).

**Figure 3.**
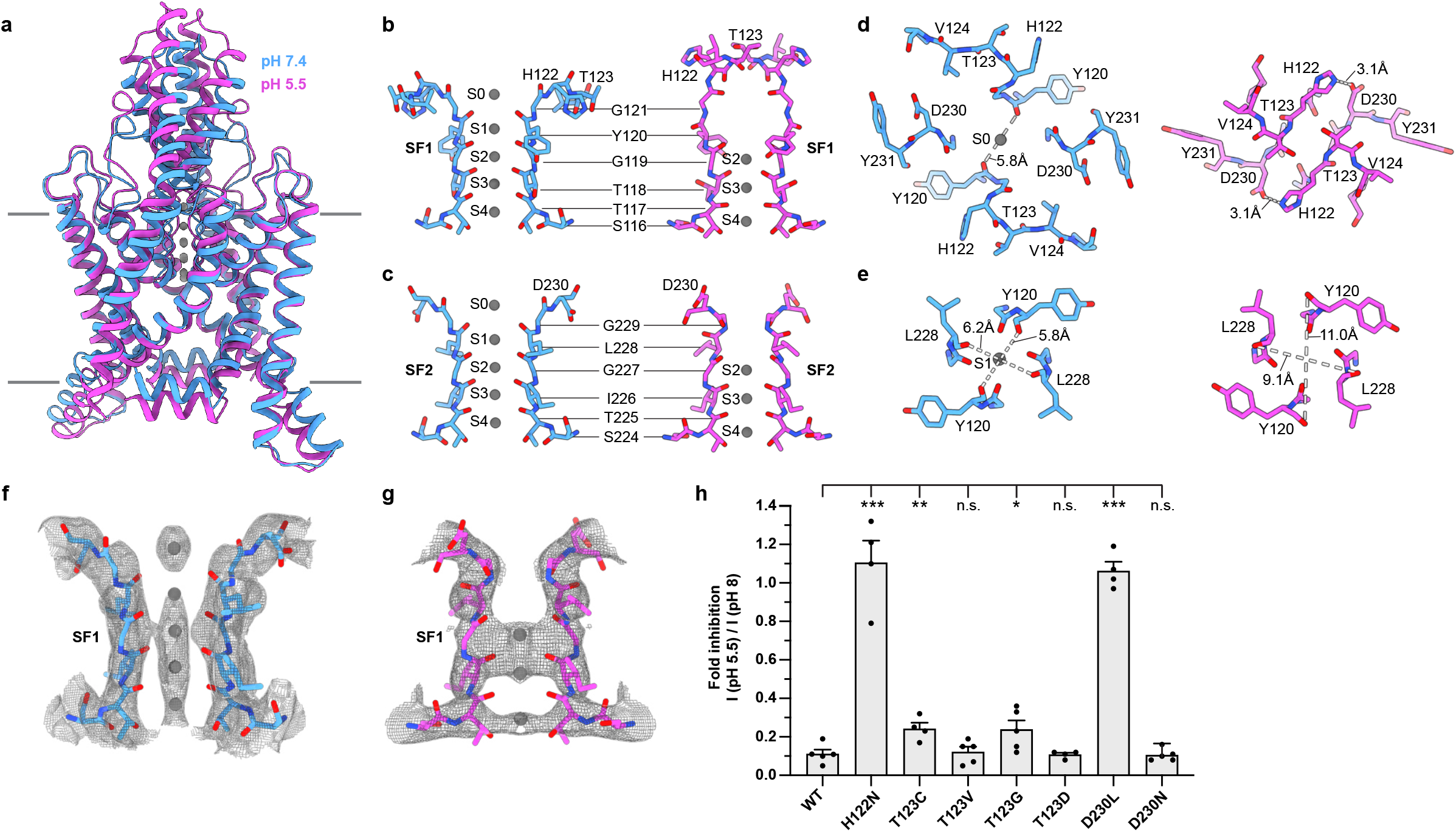
TWIK1 selectivity filter gate controlled by pH_ext_. **(a)** Overlay of open (pH 7.4, blue) and closed (pH 5.5, magenta) conformations of TWIK1 viewed from the membrane plane. **(b-e)** Comparisons of regions involved in the selectivity filter gate from open (left) and closed (right) TWIK1 structures. **(b)** SF1, **(c)** SF2, **(d)** the top of the selectivity filter, and **(e)** coordination site S1 are shown **(b**,**c)** from the membrane plane and **(d**,**e)** from the extracellular side. K^+^ coordination sites S0-S4 and intra-carbonyl or H122-D230 salt bridge distances are indicated. **(f**,**g)** Cryo-EM maps around the selectivity filter illustrating ion occupancy in **(f)** open and **(g)** closed structures. **(h)** Normalized fold inhibition of current by acidic pH_ext_ (pH_ext_ = 5.5 / pH_ext_ = 8.0 at 60 mV) for wild-type TWIK1 (0.11 ± 0.02) and mutants H122N (1.11 ± 0.22), T123C (0.24 ± 0.03), T123V (0.12 ± 0.02), T123G (0.24 ± 0.04), T123D (0.11 ± 0.01), D230L (1.09 ± 0.04), and D230N (0.13 ± 0.03). Mean plus s.e.m. were plotted for n = 5,4,4,5,5,5,4,5 cells, respectively. Differences were assessed with one-way analysis of variance (ANOVA) with Dunnett correction for multiple comparisons. ***P = < 0.0004 for H122N, D230L. **P < 0.01 for T123C. *P < 0.05 for T123G. P = 0.77, 0.88 and 0.80 for T123V, T123D, and D230N respectively (n.s.: not significant).

To test this structural model for pH_ext_ gating in TWIK1, we mutated residues implicated in pH-gating and recorded channel activity in whole cell two-electrode voltage clamp recordings. Wild-type TWIK1 is ∼90% inhibited at pH 5.5 relative to pH 8. Consistent with previous reports and its role as a proton sensor (Rajan et al, 2005), mutation of H122N abolished pH sensitivity. While T123 is integral to the pH gate, we predicted mutations at this site would have little effect on gating. This is because association of T123s at the center of the zipper above the selectivity filter at pH 5.5 is mediated by interactions between peptide backbones and the threonine side chain is exposed to solution in both channel conformations. Indeed, mutants at this site showed indistinguishable (T123V and T123D) or modestly reduced (∼75% for T123C and T123G) pH inhibition relative to the wild-type channel (despite larger current magnitudes in T123G (Fig. S5)). In contrast, replacing the acidic D230 with a hydrophobic group in D230L, which we predicted would disfavor the low pH conformation due to its proximity to protonated H122, eliminates pH inhibition. A more conservative mutation at this position, D230N, had no effect (Fig 3H). We conclude pH-gating critically involves residues in both SF1-TM2 and SF2-TM4 linkers to sense protons and seal the channel gate.

Movement of the SF1-TM2 and SF2-TM4 linkers upon protonation of H122 results in two additional large scale conformational changes. First, the entire helical cap is displaced upward ∼4 Å at low pH (Fig 4A-4B, Supplemental Video 2). Upward movement of the cap is a direct result of H122 and the SF1-TM2 linker flipping up above the selectivity filter. In its pH 5.5 position, H122-L126 clashes with E84-G89 at the bottom of EC2 (Fig. 4C). The essentially rigid body movement of EC2 and the outer half of EC1 (r.m.s.d. of L56-N95 = 0.8 Å) is made possible by P47, which kinks the otherwise continuous helix that forms TM1 and EC1. The kink straightens at low pH by approximately 12°, lifting EC1 and pushing TM1 inwards towards PH1. PH1 then bends up at its extracellular end and creates enough slack in the linker connected to EC2 for the cap to rise (Fig. 4C, S6A). The correlated cap, SF1-TM2 linker, and SF2-TM4 linker movements seal the extracellular ion access pathways to the mouth of the pore. At high pH, a wide-open path ∼3Å in radius leads from the extracellular solution to the selectivity filter above H122 and D230 and below EC2s. At low pH, the rearranged linkers form the base of a wall topped by the displaced helical cap to block ion access to the filter. While some crystal structures of K2Ps display bending of the helical cap slightly off of the two-fold symmetry axis (likely as a result of crystal contacts that necessitate this conformation) (Brohawn et al, 2012; Brohawn et al, 2013; Brohawn et al, 2019), this is, to our knowledge, the first example of helical cap involvement in K2P gating.

**Figure 4.**
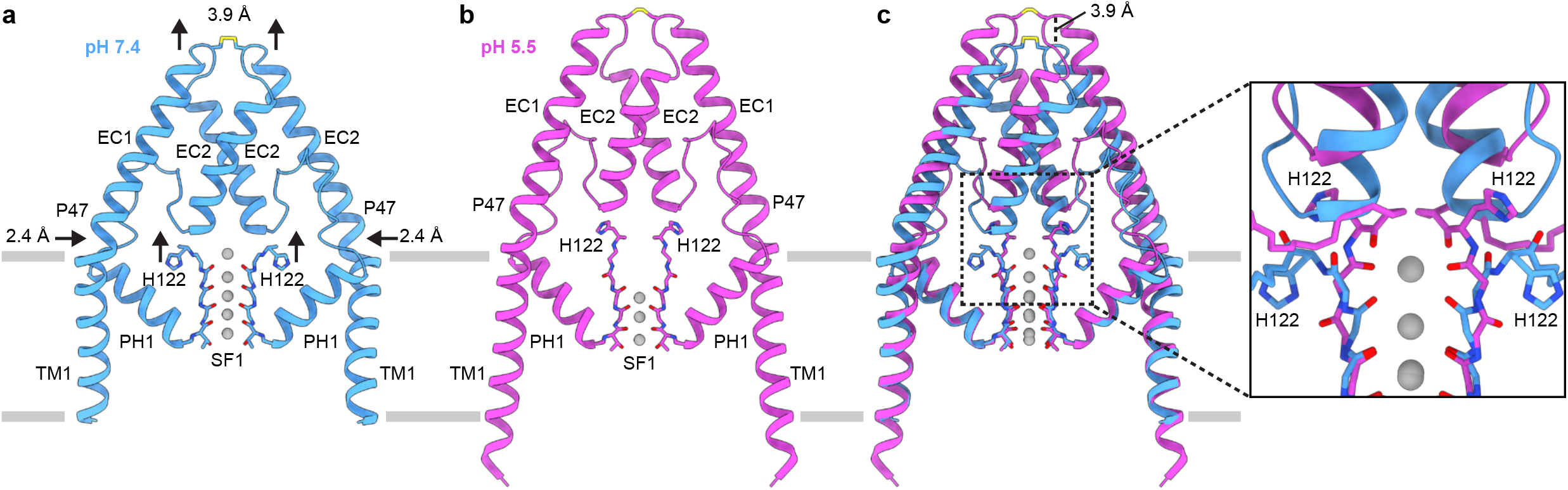
Helical cap and transmembrane helix rearrangements in response to pH_ext_. **(a-c)** View of TM1s, helical cap, PH1s, and SF1s at **(a)** pH 7.4, (**b**) pH 5.5, and **(c)** overlaid. Arrows highlight the upward movements of H122 and helical cap and the inward movement of TM1 in response to low pH_ext_. (**c, inset**) zoomed in view of region boxed in **(c)** highlighting the clash that would occur between H122 in the low pH conformation and helical cap in the high pH conformation.

**Figure 5.**
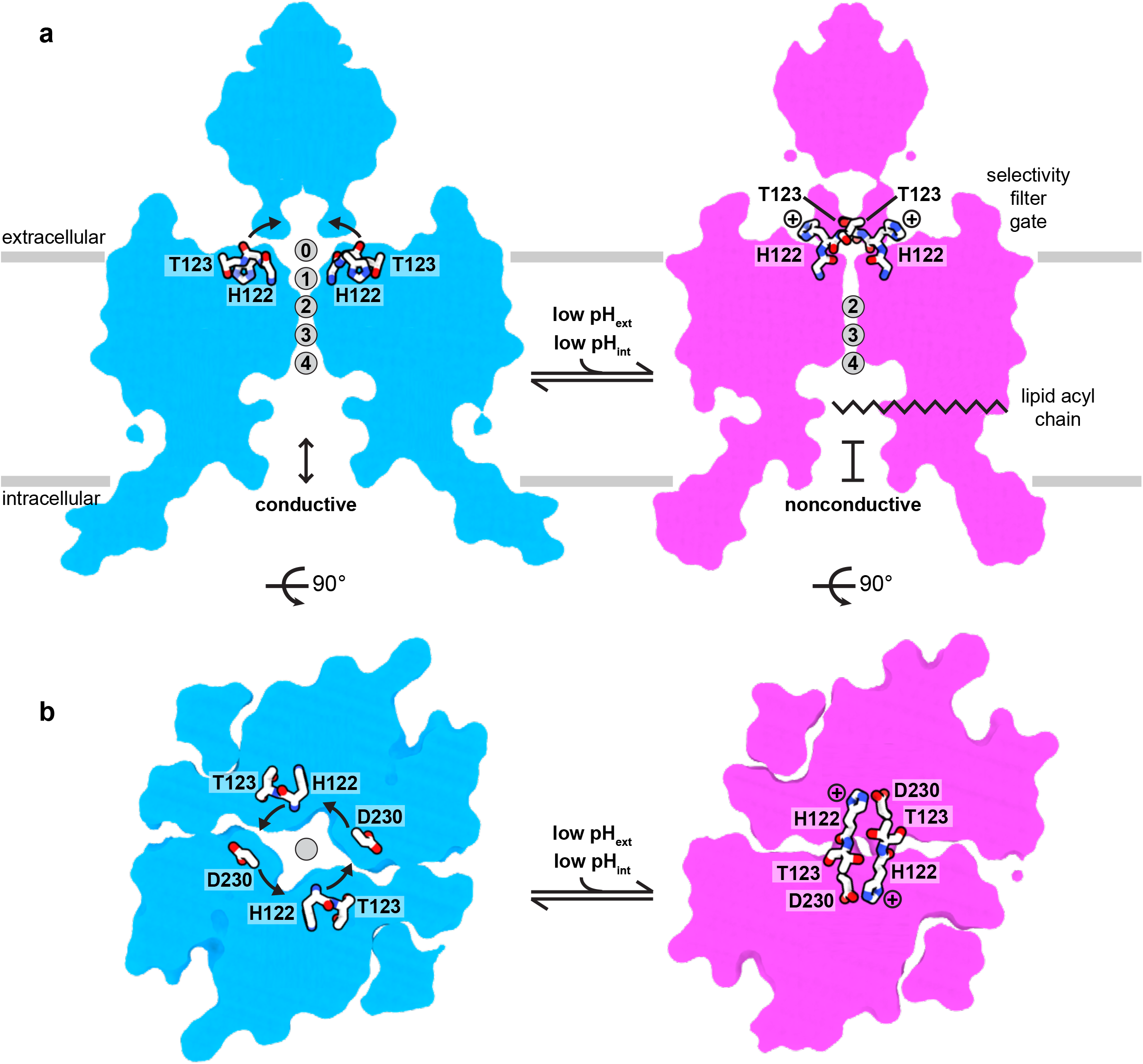
Structural model for pH gating of the TWIK1 channel. **(a**,**b)** Cartoon model for selectivity filter gating of TWIK1 gating by pH_ext_ viewed from **(a)** the membrane plane and **(b)** the extracellular side. TWIK1 is conductive at a high pH. At low pH, H122 protonation results in conformational changes that disrupt K^+^ coordination sites S0 and S1, seal the top of the selectivity filter, displace the helical cap to block extracellular ion access pathways, and dilate a lateral membrane opening to permit acyl chain binding to the channel cavity.

Second, movements of TWIK1 at low pH are propagated to the intracellular side of the membrane to dilate a lateral membrane opening, permitting acyl chain binding in the channel cavity to block conduction. TM2 rocks approximately 12° so that its intracellular half moves down towards the cytoplasm ∼2-4 Å. This is due to the top of TM2 being pulled (∼3Å by the SF1-TM2 linker rearrangement) and pushed (by straightening of TM1) towards the conduction axis (Fig. S6B). TM2 movement propagates to TM4, which tilts downwards by 4° and 1-3 Å (Fig. S6C). The resulting rearrangement of amino acids on TM2 and TM4, including I142, L146, M260, and L264, widens the lateral membrane opening from a cross sectional area of 11.8 Å^2^ at pH 7.4 to 22.6 Å^2^ at pH 5.5 at its most constricted point (Fig S7A-D). At low pH, clear tube-shaped density for a lipid acyl chain is observed extending from the dilated lateral membrane opening to just under the selectivity filter where it would sterically block ion conduction (Fig. S7C,S7G). The constriction in the lateral membrane opening at high pH is slightly smaller than the cross section of a lipid acyl-chain methylene (12.6 Å) and, consistently, similar acyl chain density is not observed in the TWIK cavity at high pH (Fig. S7B,S7F,S7H).

We note the TWIK1 lateral membrane opening at high pH is larger than that observed in other conductive K2P structures, including TRAAK and TASK2 (Fig. S7H), and just small enough to prevent lipid acyl chain access to the cavity. While this is likely the predominant conformation adopted at high pH, we cannot definitively exclude the existence of a subpopulation of particles with a slightly larger opening containing cavity bound lipids. The large lateral membrane opening at high pH may contribute to low activity of TWIK1 in cells relative to other K2Ps (Lesage et et al, 1996), since only subtle rearrangements would open it sufficiently for acyl chain binding and channel block. Indeed, the crystal structure of TWIK1 at pH 8 (Miller et al, 2012) showed a wider lateral membrane opening with acyl chains bound in the cavity and TM2, TM4, and the helical cap positioned more like the low pH than high pH structure reported here(Fig. S8), consistent with a nonconductive conformation sampled by TWIK1 at high pH and captured by crystal contacts or promoted in detergent micelles. Consistent with the notion that lipid acyl chain binding can block conduction, mutations that increase hydrophilicity of the TWIK1 cavity (L146D and L146N) around the lateral membrane opening dramatically increase channel activity (Chatelain et al 2012; Ben Soussia et al 2019). Analogous mutations activate other K2Ps; and in TRAAK, structural and single channel analyses support a model in which they disfavor lipid block and promote a TM4-down, short duration, and low conductance open state that underlies basal “leak” activity (Rietmeijer et al, 2021). Future study is necessary to determine the degree of interaction between the extracellular gate at the selectivity filter and lipid block through the lateral membrane opening in TWIK1.

Selectivity filter gating (sometimes referred to as C-type gating) has been long appreciated to play a central role in the function of many K^+^ channels (Hoshi et al, 1991), but its structural basis is just beginning to be understood. Here, we demonstrate TWIK1 utilizes a previously unobserved type of selectivity filter gate in which concerted movement of filter-adjacent linkers disrupts K^+^ coordination sites S0 and S1 and zippers shut the mouth of the channel pore. Still, one aspect of this mechanism, the upward movement of H122 and D230 above the selectivity filter in response to low pH_ext_, is reminiscent of upward movements by analogous residues in the K2P TREK1, the archaeal K^+^ channel KcsA, and the voltage-gated K^+^ channel Shaker (Fig. S9, S10). In a KcsA E71A mutant, D80 flips upward and is exposed to the extracellular solution like TWIK1 H122, but this conformational change does not block ion conduction (Cordero-Morales et al 2006) and may be promoted by interactions with an antibody Fab fragment used to crystallize the channel. In TREK1, N147 flips up in response to changes in [K^+^]_ext_, although this conformational change is only observed within one subunit and does not fully block the conduction path (Lolicato et al, 2020). Recently, a preprint described a similar upwards movement of this conserved aspartic acid (D447) in a rapidly inactivating mutant of Shaker (Tan et al, 2021). The proton sensor H122 is conserved in TRESK and pH-sensitive TASK subfamily K2Ps, and TASK1 and TASK3 have been reported to lose selectivity when exposed to low pH_ext_ like TWIK1 (Ma et al, 2012). Whether conformational changes similar to those reported here are involved in TASK channel gating and whether [K^+^]_ext_-dependent changes in selectivity are conserved between TWIK and TASK channels remains to be determined.

Other structurally characterized selectivity filter gates are more distinct from what we observe in TWIK1. The TALK-subfamily K2P TASK2, like TWIK1, is inhibited by external protons through a filter gate, but the molecular mechanisms involved are unrelated. The extracellular proton sensor of TASK2 is located further from the pore near the top of TM4 and TASK2 conformational changes at the filter are more subtle, involving rearrangement of coordination sites S0 and S1 that render the channel nonconductive (Li et al, 2020). Finally, C-type gating at the selectivity filter in an inactivating mutant of a chimeric Kv1.2-2.1 channel involves only constriction of carbonyls around S1 to disrupt K^+^ coordination (Pau et al, 2017). The discovery of a new gating mechanism in TWIK1 adds to our understanding of the myriad ways in which the selectivity filter can gate K^+^ channels to control their activity.

**Supplemental Video 1**

Extracellular view of the SF TWIK1 morphing from a conductive, high pH conformation to a non-conductive, low pH conformation.

**Supplemental Video 2**

View of TWIK1 from the plane of the membrane morphing from a conductive, high pH conformation to a non-conductive, low pH conformation.

## Methods

### Cloning, expression, and purification

Cloning, expression, and purification were performed similarly to that described for the K2P channel TRAAK^30^. A gene encoding *Rattus norvegicus* TWIK1 (Uniprot Q9Z2T2) was codon-optimized for expression in *Pichia pastoris*, synthesized (Genewiz, Inc), and cloned into a modified pPICZ-B vector (Life Technologies Inc). The resulting construct encoded a human rhinovirus 3C protease-cleavable C-terminal EGFP-10x histidine fusion protein. The resulting construct, TWIK1_-SNS-LEVLFQ/GP-(EGFP)-HHHHHHHHHH_ was used for structural studies and is referred to as TWIK1 in the text for simplicity.

Pme1 linearized pPICZ-B plasmid was transformed into *Pichia pastoris* strain SMD1163 by electroporation and transformants were selected on YPDS plates with 1mg/mL zeocin. Expression levels of individual transformants were analyzed by florescence size. For large-scale expression, overnight cultures of cells in YPD + 0.5 mg/mL Zeocin were added to BMGY to a starting OD of 1, and grown overnight at 30°C to a final OD_600_ of 25. Cells were centrifuged at 8,000 x g for 10 minutes, and added to BMMY at 27°C to induce protein expression. Expression continued for approximately 24 hours.

Cells were pelleted, flash-frozen in liquid nitrogen, and stored at −80°C. 60g of cells were broken by milling (Retsch model MM301) for 5 cycles of 3 minutes at 25 Hz. All subsequent purification steps were carried out at 4°C. Cell powder was added to 200 mL lysis/extraction buffer (50 mM Tris pH 8.0, 150 mM KCl, 1mM phenylmethysulfonyl fluoride, 1 mM EDTA, 10 µl Benzonase Nuclease (EMD milipore), 1 µM AEBSF, 1 mM E64, 1mg/ml Pepstatin A, 10 mg/ml Soy Trypsin Inhibitor, 1 mM Benzimidine, 1 mg/ml Aprotinin, 1 mg/ml Leupeptin, 1% n-Dodecyl-b-D-Maltopyranoside (DDM, Anatrace, Maumee, OH), 0.2% Cholesterol Hemisuccinate Tris Salt (CHS, Anatrace)) and then gently stirred at 4°C for 3 hours. The extraction was then centrifuged at 33,000 x g for 45 minutes. 10 ml Sepharose resin coupled to anti-GFP nanobody was added to the supernatant and stirred gently for 1 hours at 4°C. The resin was collected in a column and washed with 50 mL Buffer 1 (20 mM Tris, 150 mM KCl, 1 mM EDTA, 1% DDM, 0.2% CHS, pH 8.0), 150 mL Buffer 2 (20 mM

Tris, 300 mM KCl, 1 mM EDTA, 1% DDM, 0.2% CHS, pH 8.0), and 100 mL of Buffer 1. PPX (∼0.5 mg) was added into the washed resin in 10 mL Buffer 1 and rocked gently overnight. Cleaved TWIK1 was eluted and concentrated to ∼9.6 mL with an Amicon Ultra spin concentrator (30 kDa cutoff, MilliporeSigma, USA). The concentrated protein was subjected to size exclusion chromatography using a Superdex 200 Increase 10/300 column (GE Healthcare, Chicago, IL) run in Buffer 3 (20 mM Tris pH 8.0, 150 mM KCl, 1 mM EDTA, 1% DDM, 0.01% CHS) on a NGC system (Bio-Rad, Hercules, CA). The peak fractions were collected and spin concentrated for reconstitution.

### Nanodisc formation

10 nmol of freshly purified TWIK1 was reconstituted into nanodiscs with a 2:1:1 DOPE:POPS:POPC lipid mixture (mol:mol, Avanti, Alabaster, Alabama) at a final molar ratio of TWIK1:MSP1D1:lipid of 1:5:250 for high pH, or a ratio of TWIK1:MSPE3D1:lipid of 1:2:200 for low pH. Lipids in chloroform were mixed, dried under argon, washed with pentane, dried under argon, and dried under vacuum overnight in the dark. Dried lipids were rehydrated in buffer containing 20 mM Tris, 150 mM KCl, 1 mM EDTA, pH 8.0 and clarified by bath sonication. DDM was added to a final concentration of 8 mM. TWIK1 was mixed with lipids and incubated at 4°C for 1 hour before addition MSPE3D1 protein. After incubation for 30 minutes at 4°C, 100 mg of Biobeads SM2 (Bio-Rad, USA) (prepared by sequential washing in methanol, water, and Buffer 4 and weighed damp following bulk liquid removal) was added and the mixture was rotated at 4°C overnight. The sample was spun down to facilitate removal of solution from the Biobeads and the reconstituted channel was further purified on a Superdex 200 increase column run in 20 mM HEPES pH 7.4, 150 mM KCl, and 1 mM EDTA (high pH) or 20 mM MES pH 5.5, 150 mM KCl, and 1 mM EDTA (low pH). The peak fractions were collected and spin concentrated (50 kDa MWCO) to 1.8 mg/mL (for high pH) or 2.5 mg/ml (for low pH) for grid preparation.

### Grid preparation

The TWIK1 nanodisc samples were centrifuged at 21,000 x g for 10 min at 4°C. A 3 µL sample was applied to holey carbon, 300 mesh R1.2/1.3 gold grids (Quantifoil, Großlöbichau, Germany) that were freshly glow discharged for 30 seconds. Sample was incubated for 5 seconds at 4°C and 100% humidity prior to blotting with Whatman #1 filter paper for 3 seconds at blot force 1 and plunge-freezing in liquid ethane cooled by liquid nitrogen using a FEI Mark IV Vitrobot (FEI / Thermo Scientific, USA).

### Cryo-EM data acquisition

Grids were clipped and transferred to a FEI Talos Arctica electron microscope operated at 200 kV. Fifty frame movies were recorded on a Gatan K3 Summit direct electron detector in super-resolution counting mode with pixel size of 0.5575 Å. The electron dose was 9.039 e^−^ Å^2^ s^−1^ and 9.155 e^−^ Å^2^ s^−^ and total dose was 49.715 e^−^ Å^2^ and 50.3523 e^−^ Å^2^ in pH 7.4 and pH 5.5 datasets, respectively. Nine movies were collected around a central hole position with image shift and defocus was varied from −0.8 to −2.0 µm through SerialEM^31^. See Table S1 for data collection statistics.

### Cryo-EM data processing

For TWIK1 in nanodiscs at pH 7.4, 3,652 micrographs were corrected for beam-induced drift using MotionCor2 in Relion 3.1 and the data was binned to 1.115 A/pixel (Zheng et al, 2017; Zivanov et al, 2019). The contrast transfer function (CTF) parameters for each micrograph were determined using CTFFIND-4.1 (Rohou and Grigorieff, 2015). For particle picking in TWIK1 at high pH, 1000 particles were picked manually and subjected to reference-free 2D classification in RELION 3.1 to generate reference for autopicking. After initial cleanup through rounds of 2D classification in Relion 3.1, the remaining particles were extracted and imported into cryoSPARC (Punjani et al, 2017). CryoSPARC was used to generate an ab initio model with 2 classes and 0 similarity with or without symmetry. Particles belonging to a class with well-defined features were further refined using Homogenous refinement.

Particle positions and angles from the final cryoSPARC2 refinement job were input into Relion 3.1 (using csparc2relion.py from the UCSF PyEM (Asarnow et al, 2019)), subjected to 3D classification (4 classes, tau 4, 7.5 degrees global sampling), and refined again to produce a 4.5 Å map. Bayesian polishing, refinement, an additional round of 3D classification (4 classes, tau 16, no angular sampling), and 3D refinement improved the map further to 3.7 Å. CtfRefinement and Particle Subtraction led to a map at 3.6 Å. Particles were then repicked with Topaz (Bepler et al, 2019), and subjected to a processing procedure similar to what is described above, leading to a map at 3.4 Å. Particles leading to this map were merged with particles leading to the best map generated before repicking with Topaz, and duplicates were removed, leading to a map at 3.5 Å. This map had less anisotropy than the one produced without Topaz, and allowed for clearer visualization of the C-terminal helix. An additional round of CTF refinement, particle subtraction to remove the contribution of the nanodisc density and subsequent 3D refinement yielded a map at 3.4 Å and used for model building.

For TWIK1 in nanodiscs at pH 5.5, 7,859 micrographs were collected. The topaz model for the high pH data set described above was used to extract ∼9.6 million particles, and subjected to a data processing procedure that was similar as above, except that in this case, 2D classification was done exclusively in cryoSPARC. Bayesian particle polishing, CTF refinement, and particle subtraction yielded a final map at 3.4 Å.

### Modeling and refinement

Cryo-EM maps were sharpened using Relion LocalRes (Ramirez-Aportela et al, 2020) in the case of TWIK1 at low pH or Phenix Density Modification in the case of TWIK1 at high pH (Terwilliger et al 2020). The initial model was built from PDB 3UKM (Miller et al, 2012) and refined into the density for TWIK1 at high pH using Phenix.real_space_refine with Ramachandran and NCS restraints (Afonine et al, 2018). In the high pH structure, the bottom of TM2 and the TM2-TM3 linker (T161-A180) are much lower in resolution, possibly due to potential conformational flexibility. For this region, the model was first built in the low pH structure, using a model predicted by AlphaFold2 (Jumper et al, 2021), refined, and then docked into the high pH structure. Molprobity (Chen et al, 2010) was used to evaluate the stereochemistry and geometry of the structure for manual adjustment in Coot (Emsley et al 2010) and refinement in Phenix. Cavity measurements were made with HOLE implemented in Coot (Smart et al, 1996). Figures were prepared using PyMOL, Chimera, and ChimeraX.

### Electrophysiology

For electrophysiology, full-length TWIK1 was cloned into a modified pGEMHE vector using Xho1 and EcoR1 restriction sites such that the mRNA transcript encodes full-length TWIK1 with an additional “SNS” at the C-terminus. Three mutations (K274E, I293A, I294A) were introduced into this construct via inverse PCR to improve trafficking of the channel to the cell membrane, remove any possibility of SUMOylation, and increase basal currents. This construct was used as the background construct for all other mutants in this study, which were also generated via inverse PCR. cRNA was transcribed from Nhe1-linearized plasmids in vitro using T7 mMessage mMachine kits, and 2.5 – 3.0 ng cRNA was injected into Stage V-VI *Xenopus laevis* oocytes extracted from anaesthetized frogs.

Currents were recorded at 25 °C using two-electrode voltage clamp (TEVC) from oocytes 2 days after mRNA injection. Pipette solution contained 3 M KCl. Bath solution contained either ND96 (96 mM NaCl, 2 mM KCl, 1.8 mM CaCl_2_, 1 mM MgCl_2_, 2.5 mM Na pyruvate, 20 mM buffer), or ND96K (2 mM NaCl, 96 mM KCl, 1.8 mM CaCl_2_, 1 mM MgCl_2_, 2.5 mM Na pyruvate, 20 mM buffer) at a pH of 8, or 5.5. HEPES was used to buffer each solution to a pH of 8, and MES was to buffer to pH 5.5. Bath volume was kept at 300 uL during recording. Eggs were initially placed in the recording chamber in ND96, with ND96K at pH 8 perfused in the bath afterwards, followed by ND96K at pH 5.5. Currents were recorded and low-pass filtered at 2 kHz using a Dagan TEV-200A amplifier and digitized at 10 kHz with a Sutter Dendrite digitizer.

## Data availability

The final maps of TWIK1 in MSP1D1 nanodiscs at pH 7.4 and in MSPE3D1 nanodiscs at pH 5.5 have been deposited to the Electron Microscopy Data Bank under accession codes 25168 and 25169. Atomic coordinates have been deposited in the PDB under IDs 7SK0 and 7SK1. Original micrograph movies have been deposited to EMPIAR under accession codes EMPIAR-XXX and EMPIAR-XXX.

## Acknowledgements

We thank Dr. Jonathan Remis, Dr. Dan Toso, and Paul Tobias at UC Berkeley for assistance with microscope setup and data collection. SGB is a New York Stem Cell Foundation-Robertson Neuroscience Investigator. This work was supported by the New York Stem Cell Foundation, NIGMS grant DP2GM123496, a McKnight Foundation Scholar Award, a Klingenstein-Simons Foundation Fellowship Award, and a Sloan Research Fellowship to SGB; as well as an NSF-GRFP to TST.

---

**Table S1.**
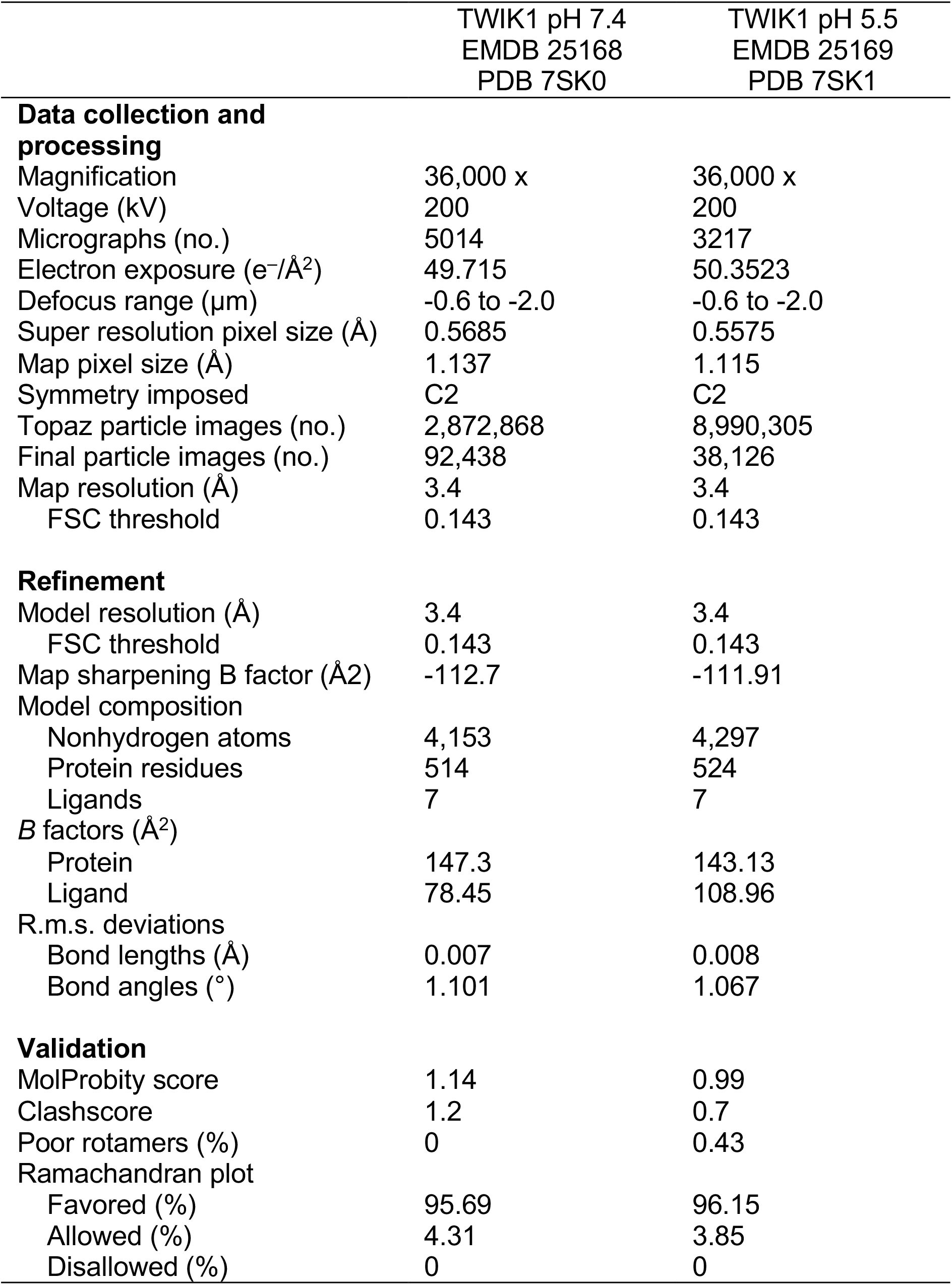
Cryo-EM data collection, refinement, and validation statistics.

**Figure S1.**
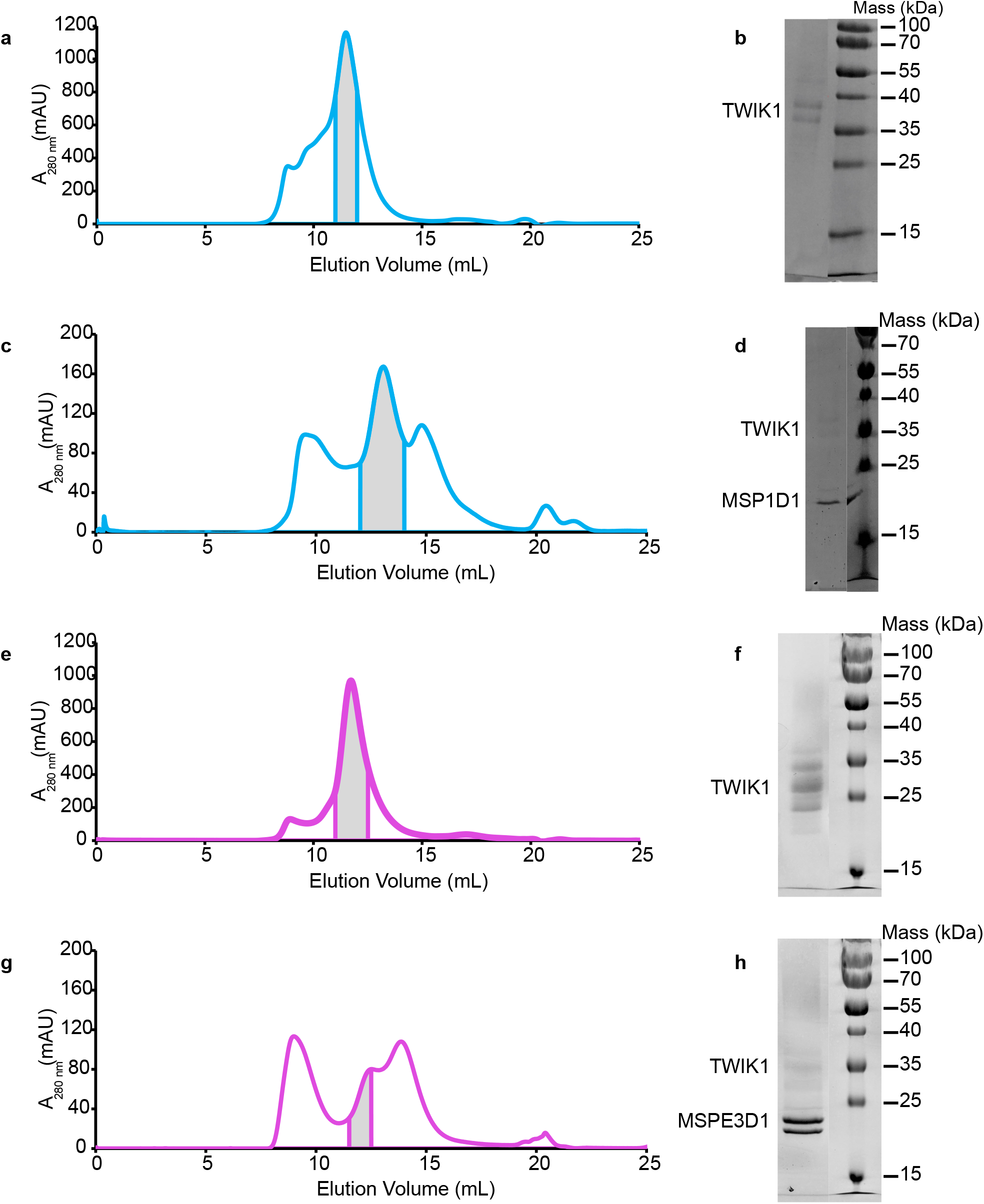
Purification and reconstitution of TWIK1. **(a-d)** Data for assembly of TWIK1-MSP1D1 nanodisc samples at pH 7.4. **(a)** Chromatogram from a Superdex 200 gel filtration of TWIK1 purified in DDM/CHS detergent. (**b**) Coomassie-stained SDS-PAGE of pooled TWIK1-containing fractions (indicated by gray bar in **(a)**). **(c)** Chromatogram from Superdex 200 gel filtration of TWIK1 reconstituted in MSP1D1 lipid nanodiscs. **(d)** Coomassie-stained SDS-PAGE of final pooled TWIK1-MSP1D1 nanodisc sample (indicated by gray bar in **(c)**). **(e-h)**, Same as **(a**-**d)**, but for TWIK1-MSPE3D1 nanodisc samples at pH 5.5.

**Figure S2.**
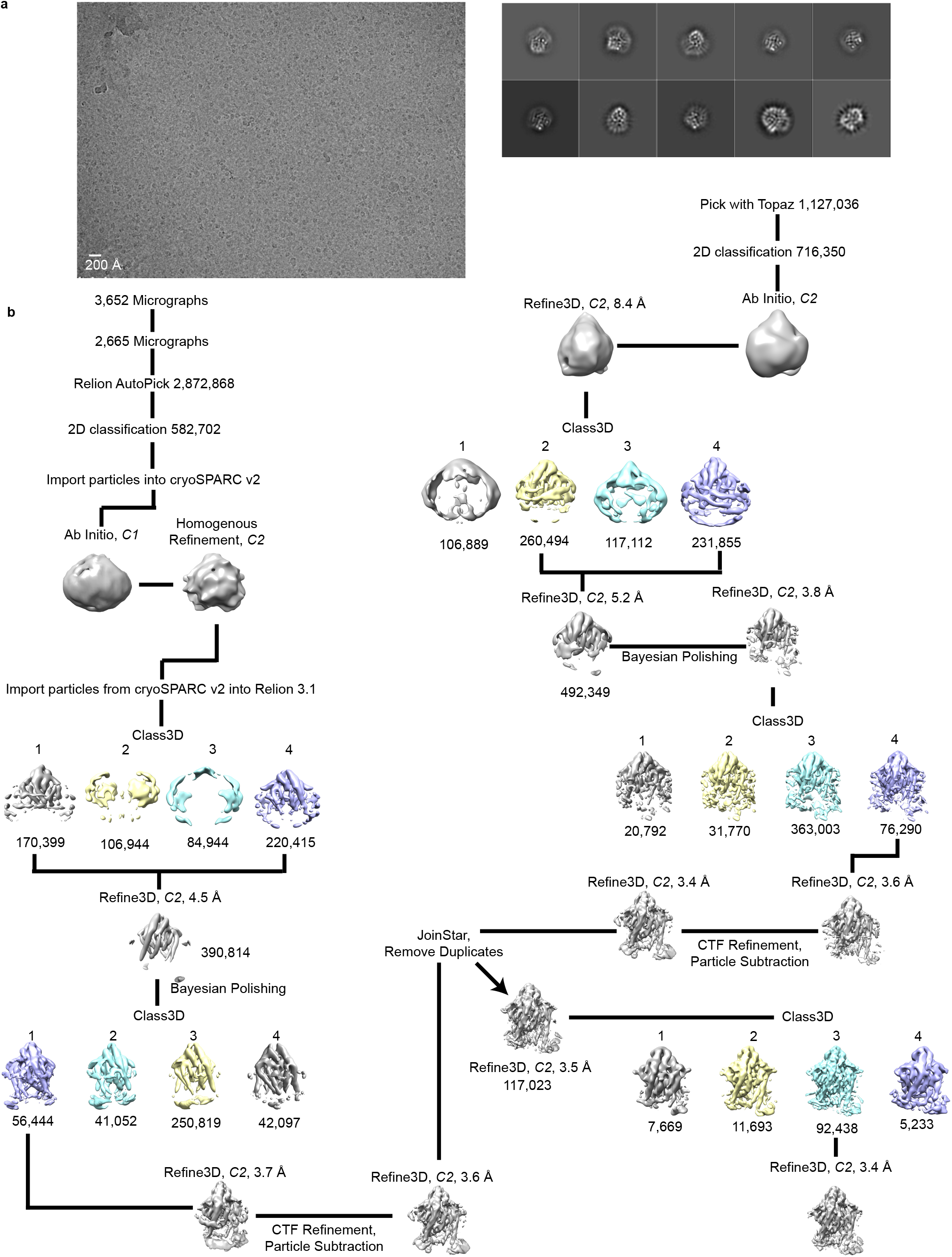
Cryo-EM processing pipeline for TWIK1 at pH 7.4. **(a)** Example micrograph (left) and selected 2D class averages (right) of TWIK1 in MSP1D1 nanodiscs at pH 7.4. 2D classification was performed in Relion with an extracted box size of 160 pix. **(b)** cryo-EM data processing steps in Relion and cryoSPARC2. See Methods for details.

**Figure S3.**
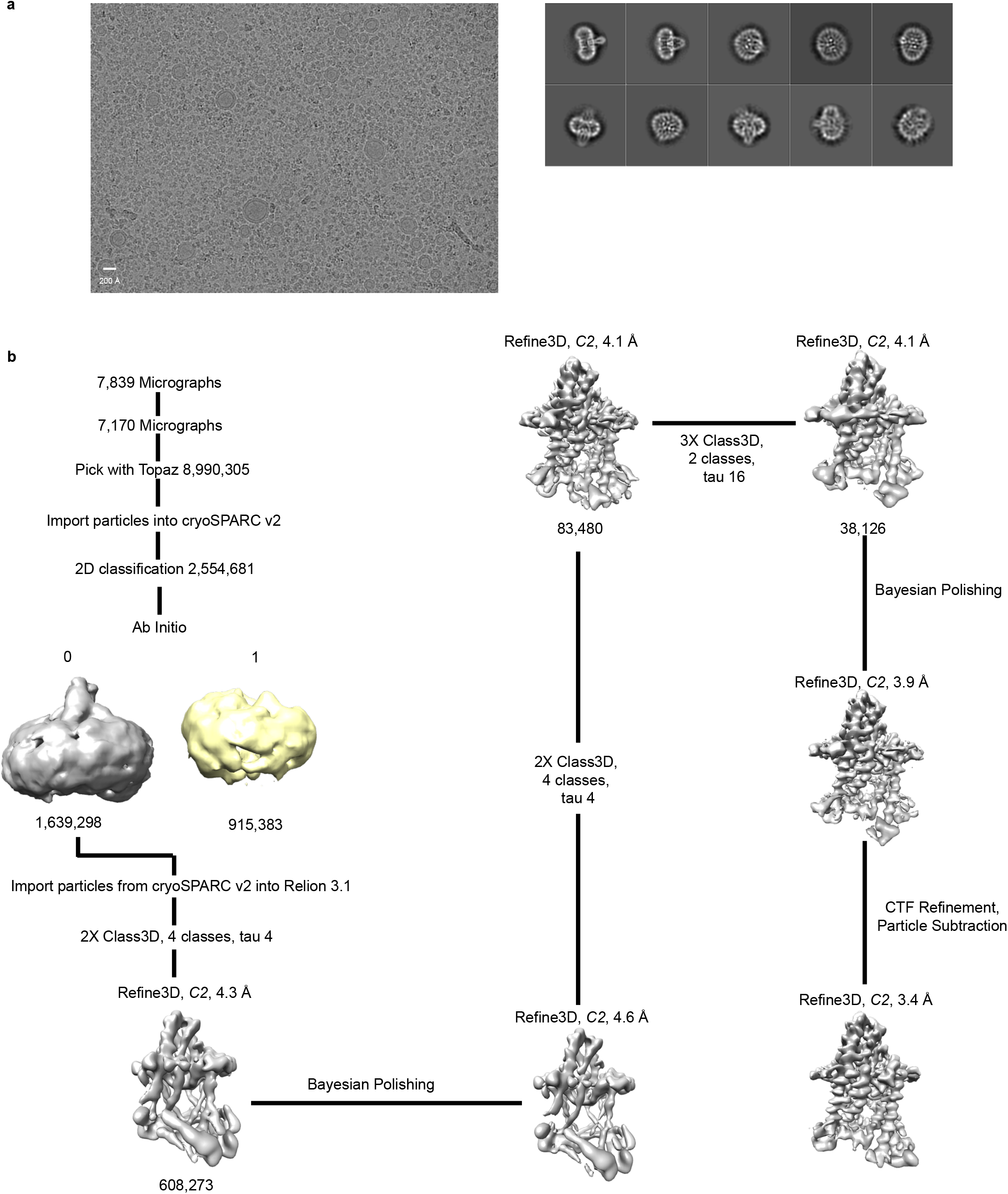
Cryo-EM processing pipeline for TWIK1 at pH 5.5. **(a)** Example micrograph (left) and selected 2D class averages (right) of TWIK1 in MSPE31D1 nanodiscs at pH 5.5. 2D classification was performed in cryoSPARC2 with an extracted box size of 200 pix. **(b)** cryo-EM data processing steps in Relion and cryoSPARC2. See Methods for details.

**Figure S4.**
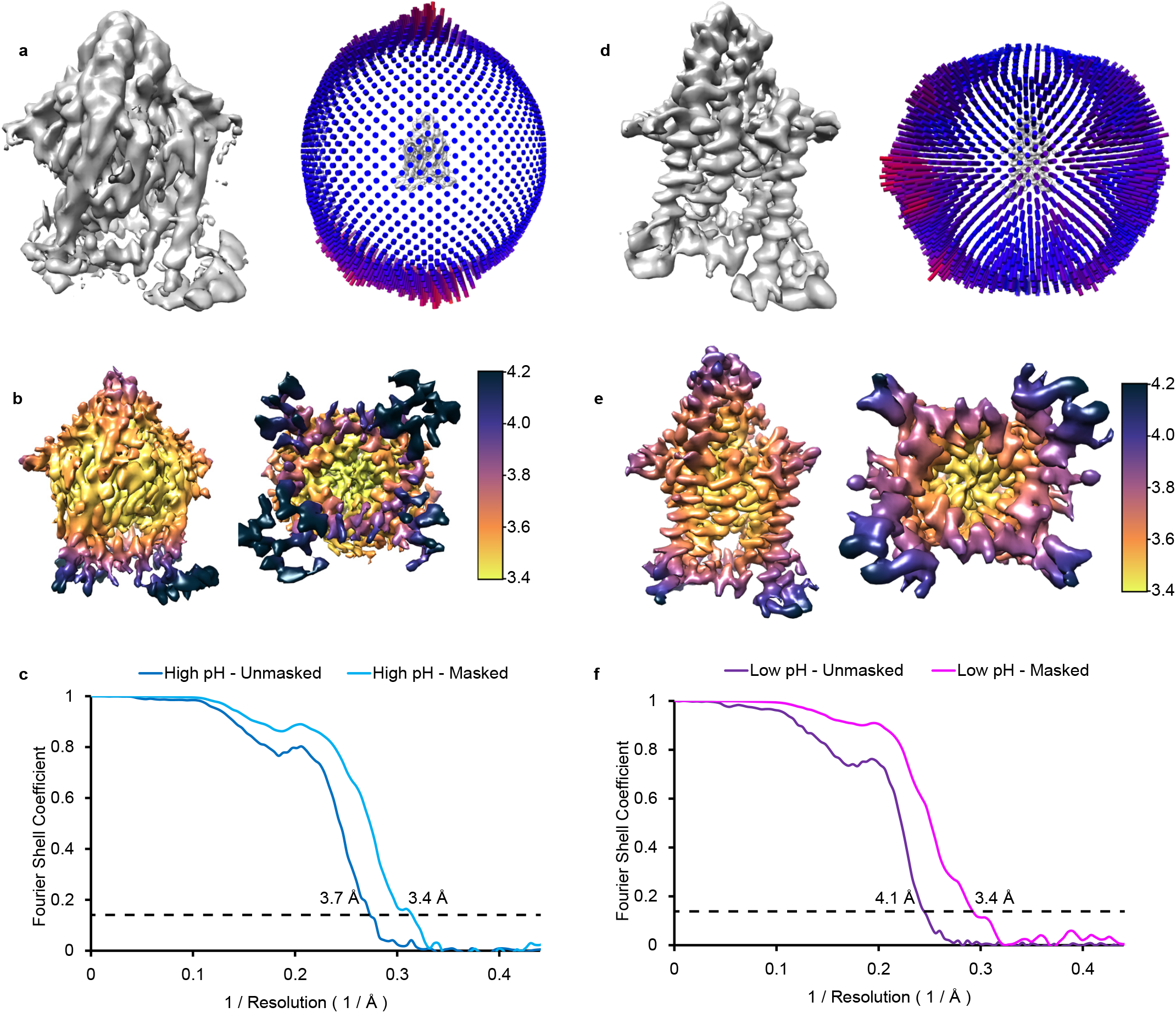
Cryo-EM Validation. Validation data for **(a-c)** pH 7.4 and **(d-f)** pH 5.5. **(a**,**d)** Angular distribution of particles used in final refinement with final map for reference. **(b**,**e)** Local resolution estimated in Relion colored as indicated on the final map. **(c**,**f)** Fourier Shell Correlation (FSC) relationships between the two unmasked or masked half-maps from refinement used for calculating overall resolution at 0.143.

**Figure S5.**
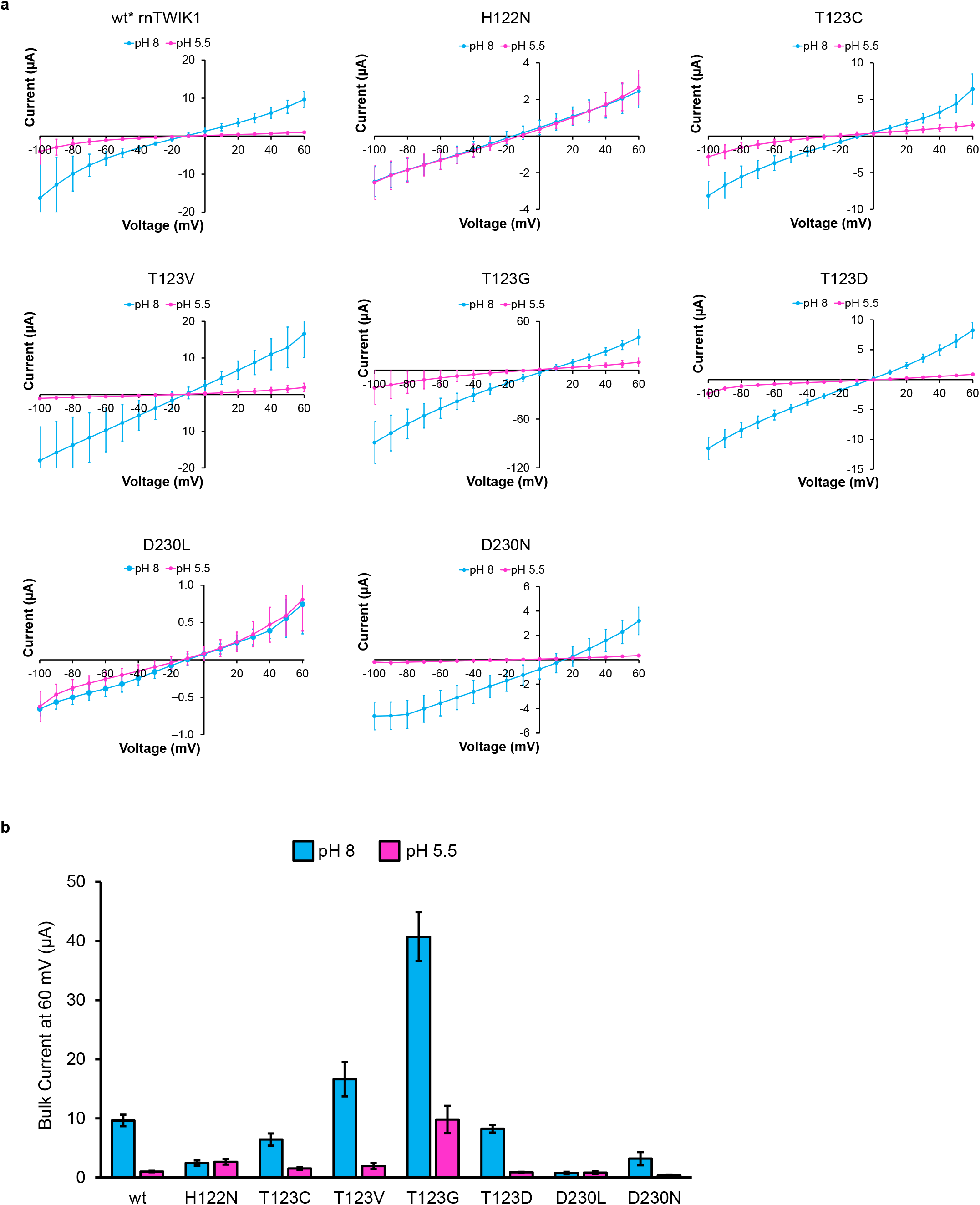
Representative IV relationships and comparison of current magnitudes for TWIK1 Mutants. **(a)** Current-voltage plot generated from currents recorded in ND96K at test potentials −100 mV to +60 mV for each TWIK1 mutant used in this study at pH 8 and pH 5.5. **(b)** Average current for each mutant at +60 mV and a pH of 8 or 5.5.

**Figure S6.**
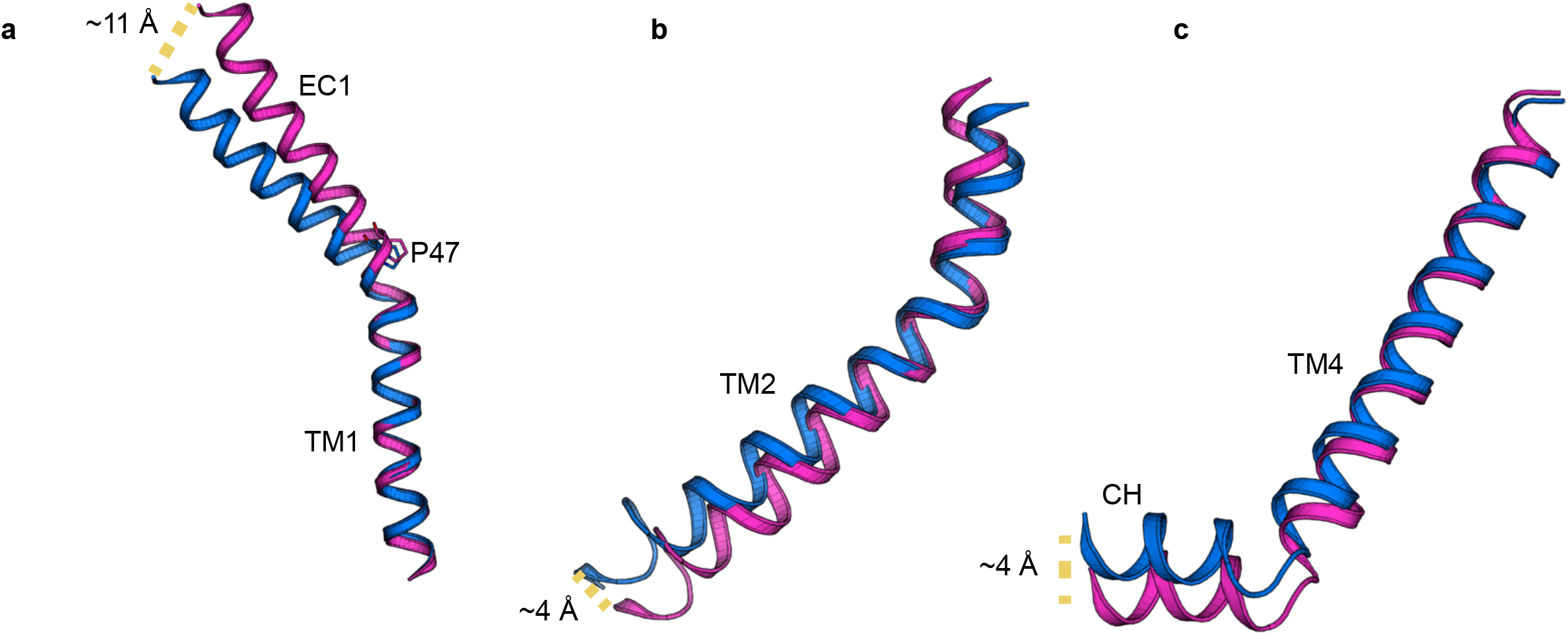
Indirect movements of TM1, TM2 and TM4 in response to pH_ext_. **(a)** Overlay of TM1 at a high pH (cyan), and low pH (magenta). **(b)** Same as in **(a)** but with TM2. **(c)** Same as in **(a)** and **(b)**, but with TM4. TM1 was aligned using residues 24-46 for each model in **(a)**. The SF and pore helices of each model were used for alignment in **(b)** and **(c)**.

**Figure S7.**
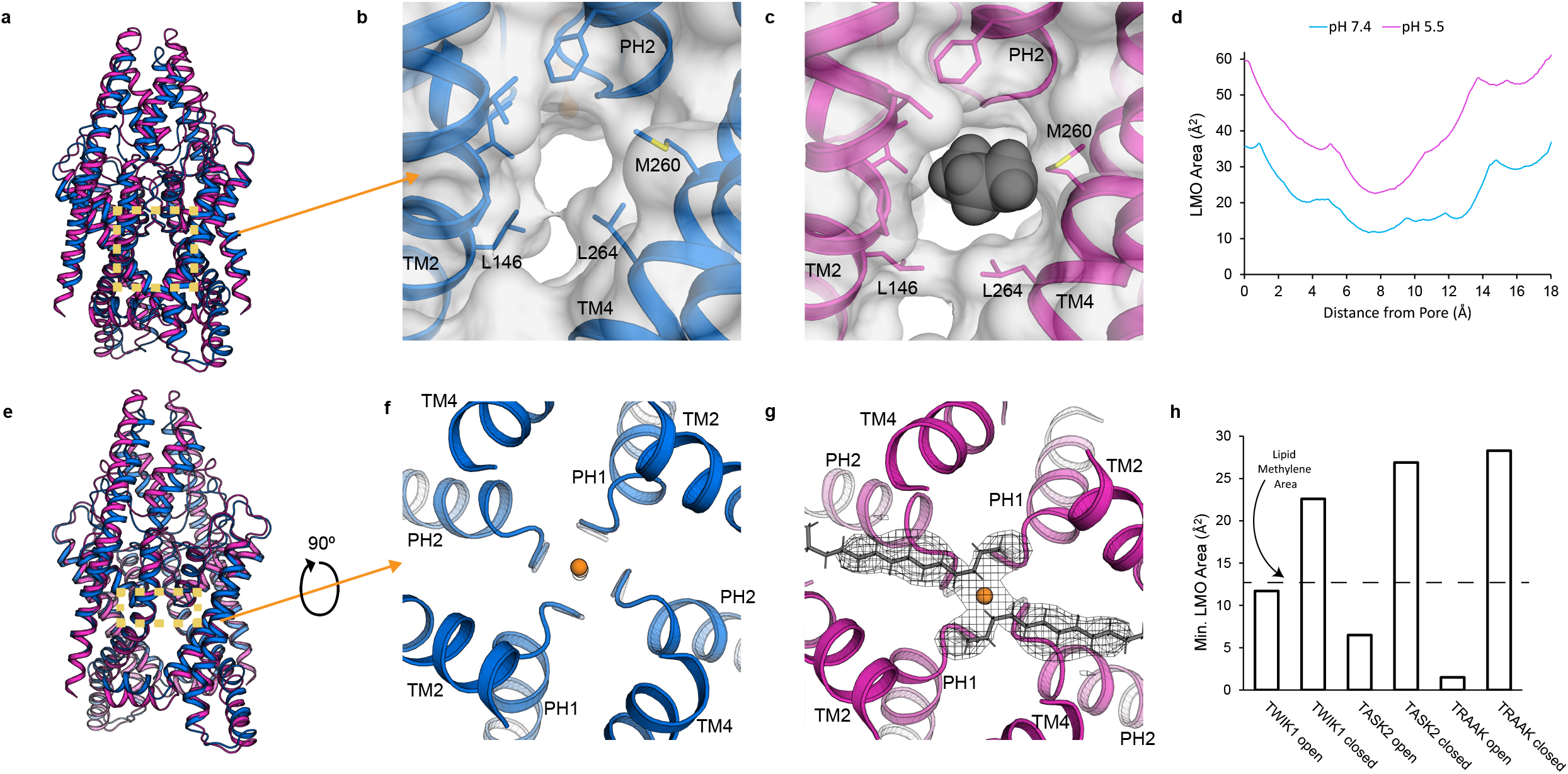
Differences in the lateral membrane opening of TWIK1 at a high and low pH. **(a)** Overlay of TWIK1 at a pH of 7.4 (blue) and 5.5 (magenta) as viewed from the plane of the membrane, with the lateral membrane opening (LMO) highlighted by a yellow box. View of the LMO at a pH of 7.4 **(b)** and a pH of 5.5 **(c). (d)** The cross-sectional area of the LMO as a function of distance from the conduction axis. **(e)** Overlay of TWIK1 at a pH of 7.4 (blue) and 5.5 (magenta) as viewed from the plane of the membrane, with the area under the SF highlighted by a yellow box. View of the selectivity filter of TWIK1 from the intracellular side at a pH of 7.4 **(f)** and a pH of 5.5 **(g). (h)** Comparison of the maximum constriction of the LMO of TWIK1 in the open and closed conformations to that of TASK2 and TRAAK. (TASK2 open = PDB: 6wm0; TASK2 closed = PDB: 6wlv; TRAAK open = PDB: 4wfe; TRAAK closed = PDB: 4wff).

**Figure S8.**
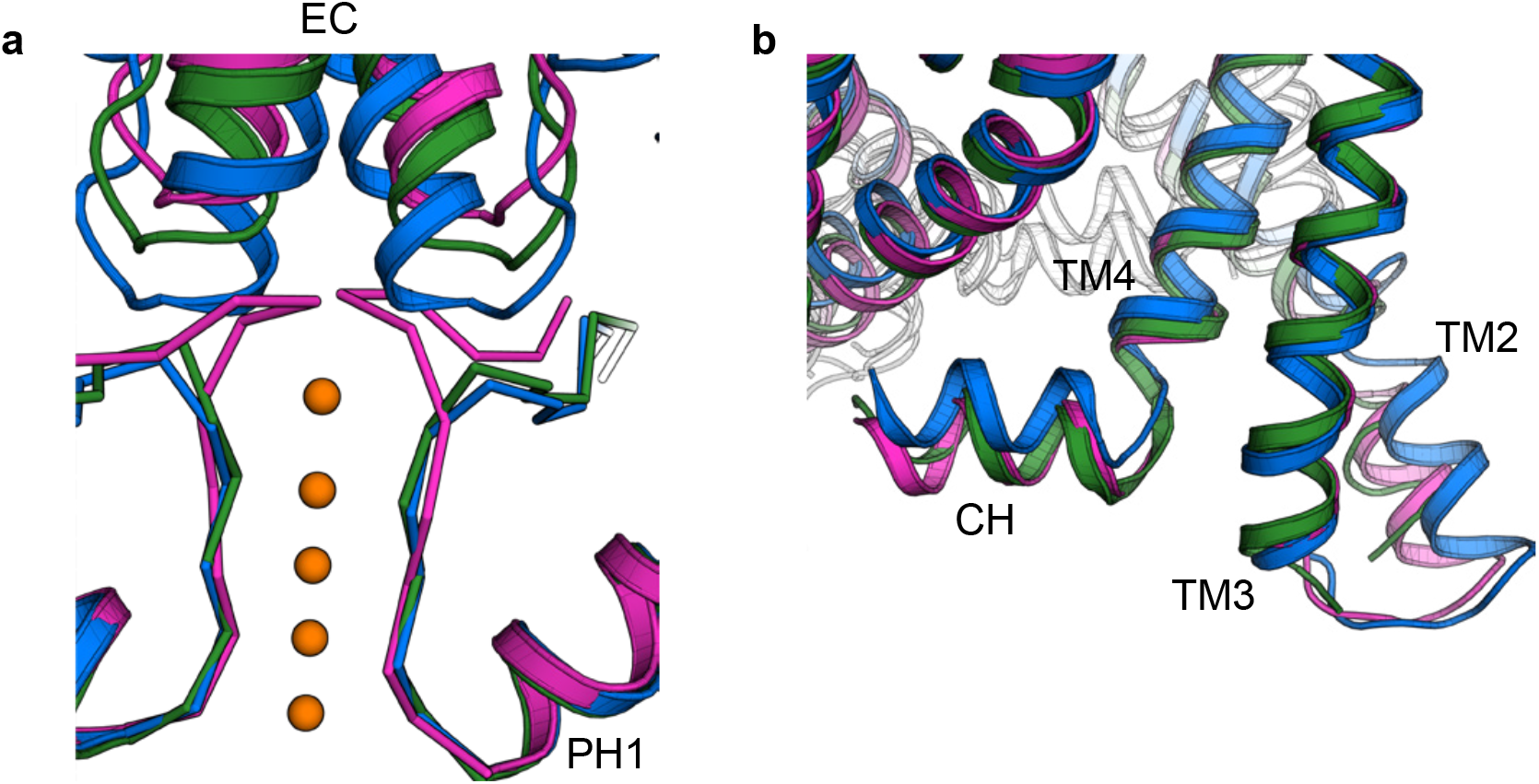
Comparison of cryo-EM structures of TWIK1 in lipid nanodiscs and a crystal structure in detergent micelles. Overlay of TWIK1 in lipid nanodiscs at a high pH (blue), at a low pH (magenta), and in detergent micelles at a high pH (green, PDB: 3ukm) focusing on the SF **(a)**, or on the intracellular ends of TM2, TM3 and TM4 **(b)**.

**Figure S9.**
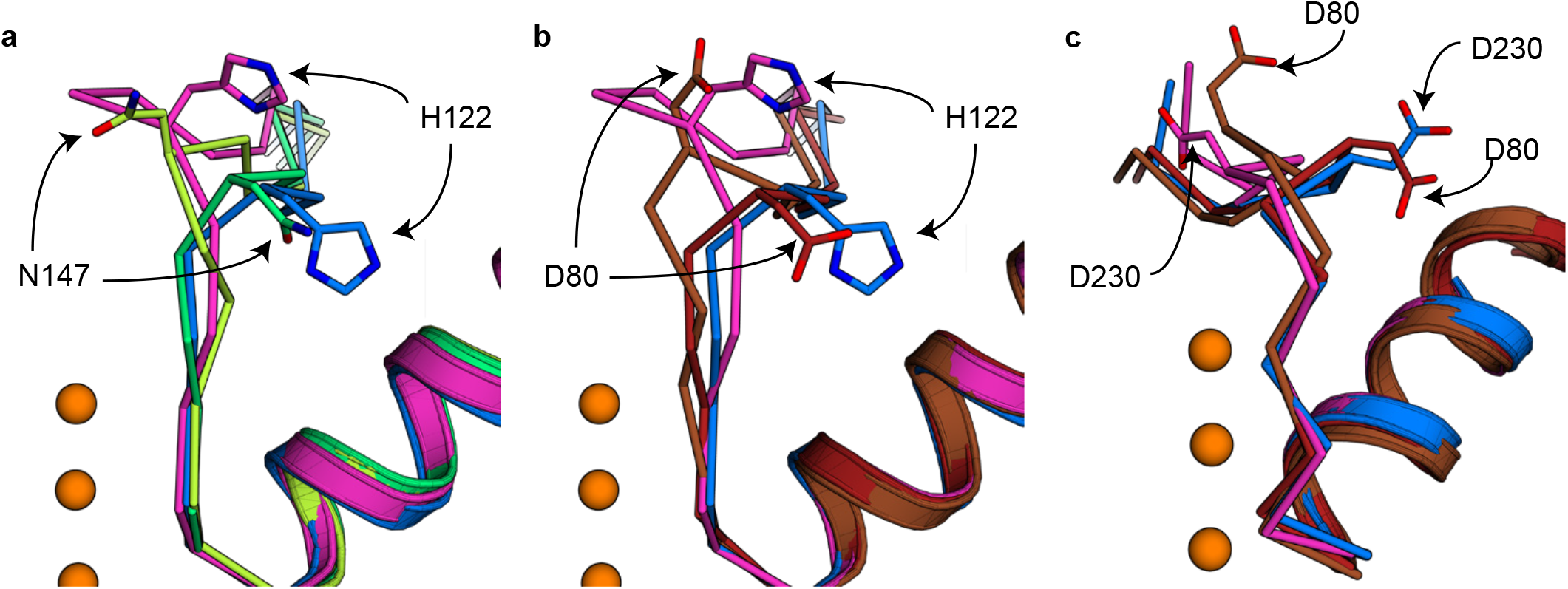
Comparison of selectivity filter gates in TWIK1 and other K^+^ channels. **(a)** View of SF1 and H122 from the plane of the membrane with TWIK1 at a low pH (magenta) high pH (blue), overlaid with TREK1 at 1 mM K^+^ (yellow-green, PDB: 6w7c) and TREK1 at 100 mM K^+^ (lime-green, PDB: 6w83). **(b)** Same as **(a)**, but with two conformations of KcsA-E71A (brown, PDB: 2atk), (dark red, PDB: 1zwi). **(c)** Same as **(b)**, but comparing SF2 and D230 of TWIK1 with KcsA-E71A.

**Figure S10.**
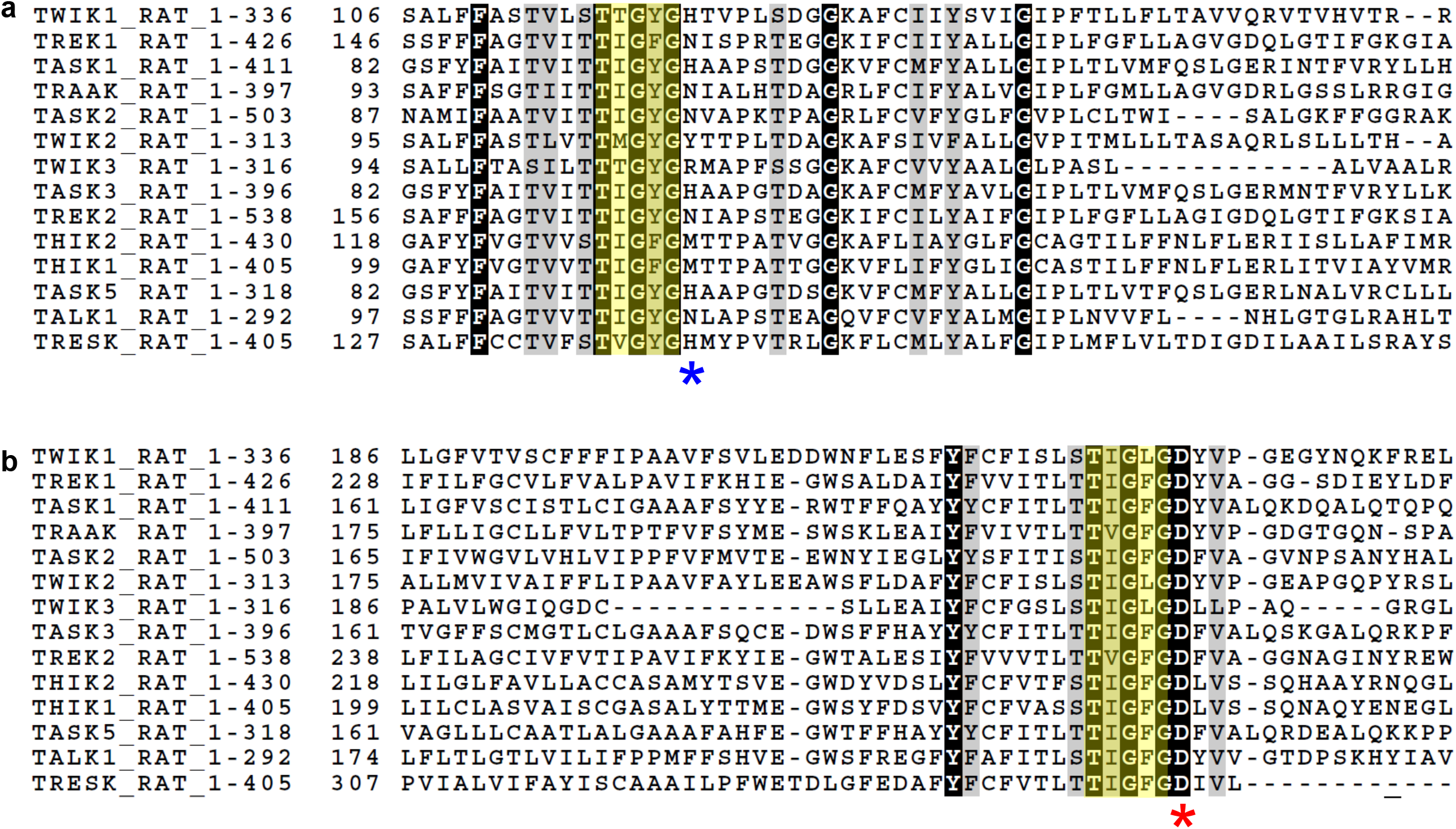
Sequence alignment of rat K2P channels. Alignment of the 14 rat K2P channels colored by conservation in a ramp from white (not conserved) to black (highly conserved), focusing on **(a)** SF1 and **(b)** SF2. The SF sequences are highlighted in yellow. Gaps in the alignment are shown as dashed black lines. H122 is highlighted with a blue star, and D230 is highlighted with a red star.

## References

Afonine, P. V. et al. Real-space refinement in PHENIX for cryo-EM and crystallography. Acta Crystallogr Sect D 74, 531–544 (2018).

Aryal, P., Abd-Wahab, F., Bucci, G., Sansom, M. S. & Tucker, S. J. Influence of lipids on the hydrophobic barrier within the pore of the TWIK-1 K2P channel. Channels 9, 44–49 (2014).

Aryal, P., Abd-Wahab, F., Bucci, G., Sansom, M. S. P. & Tucker, S. J. A hydrophobic barrier deep within the inner pore of the TWIK-1 K2P potassium channel. Nat Commun 5, 4377 (2014).

Asarnow, D., Palovcak, E., Cheng, Y. UCSF pyem v0.5. Zenodo https://doi.org/10.5281/zenodo.3576630 (2019)

Bepler, T. et al. Positive-unlabeled convolutional neural networks for particle picking in cryo-electron micrographs. Nat Methods 16, 1153–1160 (2019).

Brohawn, S. G. et al. The mechanosensitive ion channel TRAAK is localized to the mammalian node of Ranvier. Elife 8, e50403 (2019).

Brohawn, S. G., Campbell, E. B. & MacKinnon, R. Domain-swapped chain connectivity and gated membrane access in a Fab-mediated crystal of the human TRAAK K+ channel. Proc National Acad Sci 110, 2129–2134 (2013).

Brohawn, S. G., Campbell, E. B. & MacKinnon, R. Physical mechanism for gating and mechanosensitivity of the human TRAAK K+ channel. Nature 516, 126–130 (2014).

Brohawn, S. G., Mármol, J. del & MacKinnon, R. Crystal Structure of the Human K2P TRAAK, a Lipid-and Mechano-Sensitive K+ Ion Channel. Science 335, 436–441 (2012).

Chan, K. W., Sui, J.-L., Vivaudou, M. & Logothetis, D. E. Control of channel activity through a unique amino acid residue of a G protein-gated inwardly rectifying K+ channel subunit. Proc National Acad Sci 93, 14193–14198 (1996).

Chatelain, F. C. et al. TWIK1, a unique background channel with variable ion selectivity. Proc National Acad Sci 109, 5499–5504 (2012).

Chen, V. B. et al. MolProbity: all-atom structure validation for macromolecular crystallography. Acta Crystallogr Sect D Biological Crystallogr 66, 12–21 (2010).

Cohen, A., Ben-Abu, Y., Hen, S. & Zilberberg, N. A Novel Mechanism for Human K2P2.1 Channel Gating FACILITATION OF C-TYPE GATING BY PROTONATION OF EXTRACELLULAR HISTIDINE RESIDUES*. J Biol Chem 283, 19448–19455 (2008).

Cordero-Morales, J. F. et al. Molecular determinants of gating at the potassium-channel selectivity filter. Nat Struct Mol Biol 13, 311–318 (2006).

Dong, Y. Y. et al. K2P channel gating mechanisms revealed by structures of TREK-2 and a complex with Prozac. Science 347, 1256–1259 (2015).

Emsley, P., Lohkamp, B., Scott, W. G. & Cowtan, K. Features and development of Coot. Acta Crystallogr Sect D Biological Crystallogr 66, 486–501 (2010).

Enyedi, P. & Czirják, G. Molecular Background of Leak K+ Currents: Two-Pore Domain Potassium Channels. Physiol Rev 90, 559–605 (2010).

Feliciangeli, S. et al. Does Sumoylation Control K2P1/TWIK1 Background K+ Channels? Cell 130, 563–569 (2007).

Feliciangeli, S. et al. Potassium Channel Silencing by Constitutive Endocytosis and Intracellular Sequestration. J Biol Chem 285, 4798–4805 (2010).

Hoshi, T., Zagotta, W. N. & Aldrich, R. W. Two types of inactivation in Shaker K+ channels: Effects of alterations in the carboxy-terminal region. Neuron 7, 547–556 (1991).

Jumper, J. et al. Highly accurate protein structure prediction with AlphaFold. Nature 596, 583–589 (2021).

Lesage, F. et al. TWIK-1, a ubiquitous human weakly inward rectifying K+ channel with a novel structure. Embo J 15, 1004–1011 (1996).

Li, B., Rietmeijer, R. A. & Brohawn, S. G. Structural basis for pH gating of the two-pore domain K+ channel TASK2. Nature 586, 457–462 (2020).

Lolicato, M. et al. K2P channel C-type gating involves asymmetric selectivity filter order-disorder transitions. Sci Adv 6, eabc9174 (2020).

Lolicato, M. et al. K2P2.1 (TREK-1)–activator complexes reveal a cryptic selectivity filter binding site. Nature 547, 364–368 (2017).

Ma, L., Zhang, X. & Chen, H. TWIK-1 Two-Pore Domain Potassium Channels Change Ion Selectivity and Conduct Inward Leak Sodium Currents in Hypokalemia. Sci Signal 4, ra37–ra37 (2011).

Ma, L., Zhang, X., Zhou, M. & Chen, H. Acid-sensitive TWIK and TASK Two-pore Domain Potassium Channels Change Ion Selectivity and Become Permeable to Sodium in Extracellular Acidification*. J Biol Chem 287, 37145–37153 (2012).

Millar, I. D. et al. Adaptive downregulation of a quinidine-sensitive cation conductance in renal principal cells of TWIK-1 knockout mice. Pflügers Archiv 453, 107–116 (2006).

Miller, A. N. & Long, S. B. Crystal Structure of the Human Two–Pore Domain Potassium Channel K2P1. Science 335, 432–436 (2012).

Natale, A. M., Deal, P. E. & Jr., D. L. M. Structural insights into the mechanisms and pharmacology of K2P potassium channels. J Mol Biol 433, 166995 (2021).

Nematian-Ardestani, E. et al. Selectivity filter instability dominates the low intrinsic activity of the TWIK-1 K2P K+ channel. J Biol Chem 295, 610–618 (2020).

Nie, X. et al. Expression and insights on function of potassium channel TWIK-1 in mouse kidney. Pflügers Archiv 451, 479–488 (2005).

Pau, V., Zhou, Y., Ramu, Y., Xu, Y. & Lu, Z. Crystal structure of an inactivated mutant mammalian voltage-gated K+ channel. Nat Struct Mol Biol 24, 857–865 (2017).

Punjani, A., Rubinstein, J. L., Fleet, D. J. & Brubaker, M. A. cryoSPARC: algorithms for rapid unsupervised cryo-EM structure determination. Nat Methods 14, 290–296 (2017).

Rajan, S., Plant, L. D., Rabin, M. L., Butler, M. H. & Goldstein, S. A. N. Sumoylation Silences the Plasma Membrane Leak K+ Channel K2P1. Cell 121, 37–47 (2005).

Ramírez-Aportela, E. et al. Automatic local resolution-based sharpening of cryo-EM maps. Bioinformatics (2019) doi:10.1093/bioinformatics/btz671.

Rietmeijer, R. A., Sorum, B., Li, B. & Brohawn, S. G. Physical basis for distinct basal and mechanically gated activity of the human K+ channel TRAAK. Neuron 109, 2902-2913.e4 (2021).

Ritchie, T. K. et al. Chapter Eleven Reconstitution of Membrane Proteins in Phospholipid Bilayer Nanodiscs. Methods Enzymol 464, 211–231 (2009).

Rödström, K. E. J. et al. A lower X-gate in TASK channels traps inhibitors within the vestibule. Nature 582, 443–447 (2020).

Rohou, A. & Grigorieff, N. CTFFIND4: Fast and accurate defocus estimation from electron micrographs. J Struct Biol 192, 216–221 (2015).

Schewe, M. et al. A Non-canonical Voltage-Sensing Mechanism Controls Gating in K2P K+ Channels. Cell 164, 937–949 (2016).

Soussia, I. B. et al. Mutation of a single residue promotes gating of vertebrate and invertebrate two-pore domain potassium channels. Nat Commun 10, 787 (2019).

Tan, X.-F. et al. Structure of the Shaker Kv channel and mechanism of slow C-type inactivation. Biorxiv 2021.09.21.461258 (2021) doi:10.1101/2021.09.21.461258.

Terwilliger, T. C., Ludtke, S. J., Read, R. J., Adams, P. D. & Afonine, P. V. Improvement of cryo-EM maps by density modification. Nat Methods 17, 923–927 (2020).

Tsukamoto, H. et al. Structural properties determining low K+ affinity of the selectivity filter in the TWIK1 K+ channel. J Biol Chem 293, 6969–6984 (2018).

Zheng, S. Q. et al. MotionCor2: anisotropic correction of beam-induced motion for improved cryo-electron microscopy. Nat Methods 14, 331–332 (2017).

Zivanov, J. et al. New tools for automated high-resolution cryo-EM structure determination in RELION-3. Elife 7, e42166 (2018).

Zivanov, J., Nakane, T. & Scheres, S. H. W. A Bayesian approach to beam-induced motion correction in cryo-EM single-particle analysis. Iucrj 6, 5–17 (2019).

